# Transcriptomic Profiles from Normal and Tumor Tissue Samples Reveal Distinct Venule Populations and Novel Tumor Endothelial Cell Markers in Breast Cancer

**DOI:** 10.1101/2025.06.23.661087

**Authors:** Kathryn N Phoenix, Vijender Singh, Patrick A. Murphy, Kevin P Claffey

## Abstract

**Background:** The breast tumor microenvironment (TME) is a complex milieu composed of many factors contributing to breast cancer (BC) heterogeneity and therapeutic resistance. Aberrant tumor vasculature in the TME limits nutrient and drug delivery, inhibits anti-tumor immunity, and contributes to a lack of cancer therapy efficacy. Utilizing publicly available scRNA-seq datasets, this study characterizes differences between normal breast and breast tumor endothelial cells (EC), provides insights into tumor endothelial cell subtypes, endothelial anergy, and identifies novel, tumor-specific vascular therapeutic targets.

**Methods:** Gene expression data from normal and breast tumor tissue samples were integrated, and the EC subset was extracted via canonical gene marker expression. The EC subset was clustered and evaluated for cell subtypes and differentially expressed genes (DEG). Normal EC (NEC) and tumor EC (TEC) markers were further assessed for correlation to bulk gene expression and patient survival outcomes in cBioPortal and Kaplan-Meier Plotter. Cell type gene expression specificity was evaluated in the 3CA single-cell RNA-seq datasets across multiple cancers.

**Results:** This analysis revealed differences in NEC and TEC subtype populations. Breast NEC contained similar proportions of venule and capillary populations, while breast TEC demonstrated a majority of the venule subtype. Further, TEC venules were phenotypically distinct from the NEC venules. Consistent with endothelial anergy, suppression of the key adhesion protein SELE was noted, as well as several pro-inflammatory cytokines including IL6, CCL2, and CXCL8, likely downstream of aberrant NF-kB signaling. Differential gene expression analysis identified several TEC specific up-regulated genes compared to NEC, including CLEC14a, IGFBP4, EMCN, and ADM5. CLEC14a, EMCN, and ADM5 were further validated in the single-cell Curated Cancer Cell Atlas (3CA) to be highly specific to the endothelial cell clusters across multiple tumor types, while IGFBP4 was diversely expressed in endothelial, fibroblast, and some malignant cell types. ADM5, a novel tumor vascular marker, was enhanced in TEC venules and less so in arteriole or capillaries. High expression of ADM5 was associated with poor breast cancer patient survival in the basal PAM50 cancer subtype compared to normal and luminal subtypes. Further, across multiple cancer types, high ADM5 expression was associated with reduced patient survival in anti-PD1– and anti-CTLA4-treated patients but not in anti-PDL-treated patients.

**Conclusions:** Integration of single-cell RNA-seq data identified an anergic-like response in breast TEC and multiple, highly specific markers to TEC not found in normal breast tissue. CLEC14a and EMCN were validated as TEC markers, extending their annotation in breast TEC, and ADM5 identified as a novel TEC marker in breast and other cancers. Moreover, as ADM5 is associated with reduced patient overall survival, this data suggests that a better understanding of ADM5 and other TEC-specific response pathways may provide novel approaches to reactivate anergic TECs and lead to effective therapeutic interventions for cancer patients.

Graphical Abstract

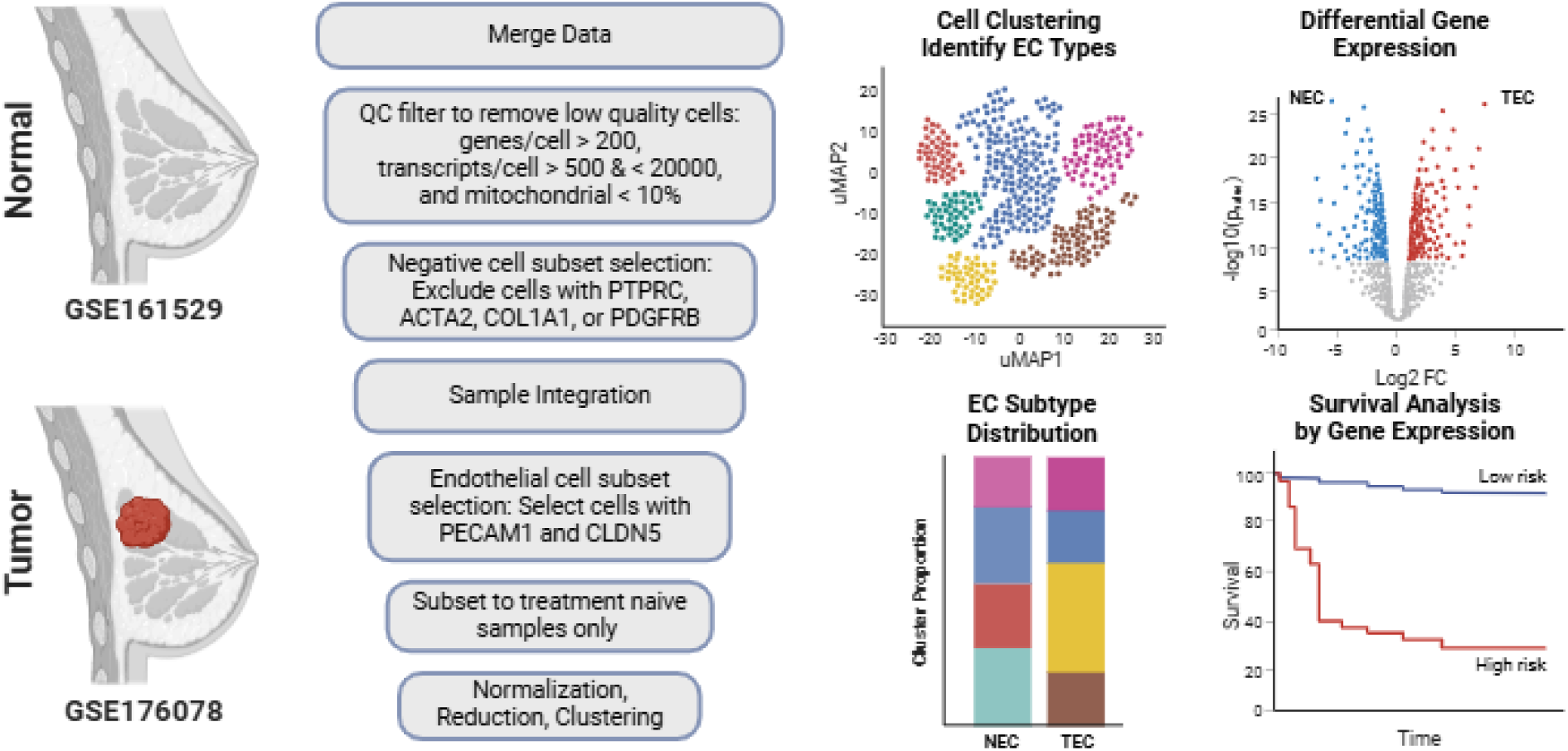

## Introduction

The tumor microenvironment (TME) is a heterogeneous and complex composition of many factors that play a role in BC heterogeneity and therapeutic resistance. The TME consists of a multitude of features, including malignant epithelial cells, stroma, immune cells, blood supply, lymphatics, and extracellular matrix (ECM), providing critical factors and interactions required for tumor progression.[1–9] As a tumor expands, blood vessels develop to provide nutrients and enable tumor growth and metastatic dissemination. However, tumor vasculature is abnormal and consists of vessels with insufficient perfusion and a disorganized, leaky vascular network.[10–13] Aberrant tumor vasculature is known to limit nutrient and drug delivery, inhibit anti-tumor immunity, and contribute to a lack of cancer therapy efficacy.[14,15] Deficient tumor blood vessel function leads to persistent regions of hypoxia and acidic pH, increased tumor heterogeneity, and impaired endothelial cell function. One consequence of this dysfunction is endothelial anergy, a phenotype that inhibits immune cell recruitment of the vasculature, suppressing adhesion molecule expression and inflammatory signaling. This immune evasion mechanism contributes to resistance against chemotherapy, radiation, and immunotherapy.[16–20] In anergenic endothelium, the response to inflammation is blunted and adhesion molecules such as ICAM1, VCAM1, and SELE are downregulated relative to normal endothelium, limiting leukocyte trafficking into tumors. Repression of inflammatory signaling further reduces immune cell recruitment, weakening anti-tumor immunity.[21] However, the mechanisms and endothelial cell (EC) phenotypic markers of vascular anergy in cancer are poorly understood and under-studied in breast cancer. To overcome tumor endothelial anergy and the resulting therapeutic resistance, novel therapies and combinatory approaches are needed.

Anti-angiogenic therapies (AAT) have shown some promise in improving therapeutic response in cancer patients when used in combination with other therapeutic approaches, including chemotherapy and immunotherapy.[22–25] Most of the current AAT target the vascular endothelial growth factor (VEGF) pathway (e.g. bevacizumab, ramucirumab, and small molecule pan-tyrosine kinase inhibitors). While some cancer patients benefit from anti-VEGF/VEGFR targeting, many cancer patients do not respond or develop resistance to anti-angiogenic therapies.[26–29] In breast cancer, the early approval of bevacizumab was retracted following several phase III clinical trials that showed no improvement in pCR, PFS, or OS along with a high number of grade III adverse events.[30] Given the limitations of VEGF-targeting therapies, expanding the therapeutic landscape to include vascular activation strategies may improve patient outcomes. Given that endothelial anergy limits immune cell recruitment and contributes to tumor immune evasion, targeting tumor endothelial cell (TEC) specific pathways that modulate endothelial activation may offer new therapeutic opportunities beyond conventional anti-angiogenic approaches. Approaches such as targeting TEC-specific metabolic adaptations, ECM remodeling, or immune-modulating endothelial pathways represent potential avenues for overcoming resistance and enhancing treatment efficacy.

EC are heterogeneous across vessel type and tissue localization and demonstrate broad phenotypic plasticity.[31–34] In tumor angiogenesis, EC are recruited from established vasculature or derived from endothelial progenitor cells in response to angiogenic factors released from the tumor.[11,35–37] These cells migrate, proliferate, and develop into tumor vasculature, becoming phenotypically distinct tumor endothelial cells (TEC). Previous work in isolated and cultured primary breast TEC demonstrated distinct differences in TEC function compared to normal breast tissue-derived EC. TEC displayed increased proliferation, migration, and angiogenic potential as well as exhibited intrinsic resistance to the anti-angiogenic small molecule sorafenib.[38] Additionally, bulk RNA-seq and single-cell RNA-seq (scRNA-seq) analyses of TEC revealed clear differences between tumor and adjacent tissue EC gene expression profiles as well as distinct subgroups of TEC within the same tumor and between various tumor types, including breast, lung, prostate, and colon.[39–42] Transcriptomic studies on breast TEC have been previously limited by the number of samples analyzed, restricted breast tumor subtypes, lack of normal tissue samples, use of pre-enriched TEC samples, and a limited depth of investigation.[39,43–45] In this work, an inclusive analysis of breast tumors and normal breast tissues was performed to understand the phenotypic differences between breast tumors and normal vascular endothelial cells. This study conducts a comprehensive scRNA-seq analysis to characterize endothelial heterogeneity in breast cancer, identifying NEC and TEC subtypes, markers of endothelial anergy, and potential vascular therapeutic targets. By leveraging publicly available datasets, this work provides insights into TEC-specific phenotypic shifts, immune modulation, and novel opportunities for therapeutic intervention.

## Methods

### Breast scRNA-seq Data Review and Selection for Analysis

A literature search was conducted for scRNA-seq analyses from breast cancer and normal breast tissue samples that had publicly available datasets from 2020 through 2023. Identified studies were reviewed for the number of patient samples, method of tissue processing, cell numbers per sample, and breast tumor subtypes included in the analyses, **Table 1**. Two studies were selected for endothelial cell population analysis: GSE176078 and GSE161529.[46,47]

**Table 1:** Summary of breast scRNA-seq studies that include vascular components.

| First Author | Year | Tissue | No of Pts | Enrichment | GEO |
| --- | --- | --- | --- | --- | --- |
| Wu | 2021 | breast tumor | 26 | No | GSE176078 |
| Pal | 2021 | normal, preneoplastic,<br>and breast tumor | 55 | No | GSE161529 |
| Tlsty | 2021 | normal breast | 14 | Yes | GSE190592 |
| Bhat-Nakshatri | 2021 | normal breast | 11 | No | GSE164898 |
| Xu | 2021 | breast tumor and<br>matched lymph nodes | 5 | No | GSE180286 |
| Geldhof | 2022 | breast tumor and<br>matched peritumoral tissue | 9 | Yes | GSE155109 |
| Gray | 2022 | normal breast | 16 | No | GSE180878 |
| Lui | 2022 | breast tumor and<br>matched lymph nodes | 8 | No | GSE167036 |
| Kumar | 2023 | normal breast | 126 | No | GSE195665 |
| Hyunsoo | 2023 | normal and breast tumor | 10 | No | GSE245601 |

### Data Integration and Seurat scRNA-seq Analysis Workflow

Data samples were merged, integrated, and analyzed in R via standard Seurat methods as previously described.[48–51] Briefly, total cells per sample were filtered for cell quality metrics including mitochondrial gene expression, number of genes per cell, and number of transcripts per cell. Cells with less than 200 genes per cell, less than 500 or greater than 20,000 transcripts per cell, and mitochondrial gene expression of more than 10% were excluded. Direct endothelial cell selection was achieved through subset selection of cell types established by canonical gene expression.

**Negative cell selection**: PTPRC – hematopoietic cells, ACTA2 – vascular smooth muscle cells, COL1A1 – fibroblasts, PDGFRB – vascular smooth muscle cells and pericytes

**Positive cell selection**: PECAM1 and CLDN5 – endothelial cells

Normalized, integrated data was scaled against all genes prior to RunPCA, RunUMAP, FindNeighbors (KNN), and FindClusters function analyses. Dimension reduction was performed using the top eleven principal components (PC) as determined by the inflection of the change in variation between consecutive PCs less than 0.1%.

### Differential Gene Expression

Differentially expressed genes (DEG) were identified using a pseudobulk method analysis of the scRNA-seq data in Seurat. Pseudobulk samples were generated with the AggregateExpression function and grouped by sample tissue source (Normal or Tumor). Following psuedobulk sample aggregation, DEG was determined by the FindMarkers function comparing all normal sample and tumor sample endothelial cells, applying the MAST package to run differential expression testing with min.pct = 0.1, and Bonferroni correction to establish adjusted p-values.[48,51,52] Genes that were not expressed (pct = 0) in either NEC or TEC were excluded from further analysis to reduce potential false positives.

### Gene Ontology and Gene Signature Enrichment Analysis

Gene Ontology analysis was performed with the enrichGO function of clusterProfiler with a q-value cut-off of 0.05 and adjusted p-value cutoff of 0.1. The top 50 clusters were plotted on an enrichment network map (emapplot) with enrichplot. Gene Signature Enrichment Analysis (GSEA) was performed using GSEA v4.2.3 software package with a pre-ranked list of genes compared to the hallmark gene sets from MSigDB.[53–55]

### Bulk RNA Expression and Correlation Analysis

cBioPortal for Cancer Genomics was utilized to evaluate bulk RNA expression data and gene expression correlation (Spearman’s correlation (r_s_) – no correlation (0-0.19), weak correlation (0.2-0.39), moderate correlation (0.4-0.59), strong correlation (0.6-0.79), very strong (0.8-1.0)).[56–58] Breast cancer bulk transcriptomic data was assessed from the METABRIC (n=2509) (https://bit.ly/45mj8Aq) and TCGA (n=1084) (https://bit.ly/3ScbAZA) studies.[59,60]

### Cell Type Specificity Analysis

The Curated Cancer Cell Atlas (3CA) online database (https://www.weizmann.ac.il/sites/3CA/)was utilized to assess the cell type-specific expression of genes across multiple cancer types and datasets.[61]

### Survival Analysis

Patient survival was assessed with Cox proportional hazards regression analysis using the Kaplan-Meier Plotter online tool with RNA-seq data from GSE96058.[62,63] (https://kmplot.com/analysis/index.php?p=service&cancer=breast_rnaseq_gse96058) Patient gene expression was divided into high– and low-expression groups and analyzed for correlation with overall survival (OS) and progression-free survival (PFS). Additional survival data was conducted in cBioPortal with the TCGA breast cancel dataset. [56–58,60]

## Results

### Data Acquisition, Processing, and Endothelial Cell Identification in Breast Tissue Samples

To compare endothelial cell profiles from normal and tumor breast tissue transcriptomic data, a review of published or publicly available single-cell RNA sequencing datasets was conducted in NCBI Gene Expression Omnibus (GEO). A total of 11 datasets were identified as available through the end of 2023. Each was evaluated for tissue source and quality, sample number, cell number, and cell processing methods, **Table 1**. To be selected for this analysis, datasets were required to be publicly available, include more than 10 patients with diverse BC subtypes, and to have analyzed entire tissue samples (e.g. no pre-enrichment), no sample pooling, primary tumors only (no metastatic tumors or lymph nodes), and only treatment naïve patient samples were included (no therapeutically treated patients or ex vivo tissue treatment). Two datasets, GSE176078 and GSE161529, were selected for further analysis. Preliminary evaluation of each dataset revealed that all tumor samples from GSE176078 and the normal samples from GSE161529 displayed the expected range of endothelial cells; however, GSE161529 tumor samples were not included due to low quality and count of endothelial cells per sample (GSE161529 average EC%/tumor sample = 0.9±1.3 vs GSE161529 average EC%/normal sample = 2.5±2.2, p=0.029; vs GSE176078 average EC%/tumor sample = 2.6±2.7, p=0.0098). Following sample data merge and integration, directed cell subset selection was used to isolate endothelial cells. Cells expressing markers of hematopoietic cells, vascular smooth muscle cells, fibroblasts, and pericytes were excluded, while cells expressing both PECAM1 and CLDN5 were selected and identified broadly as EC. A comparison of each sample type revealed no significant differences between normal and tumor total cell counts or the percentage of endothelial cells identified, **Figure 1A**. A final count of n=22 breast tumor-derived samples and n=2176 EC along with n=13 normal breast tissue-derived samples with n=1006 EC were used in the analysis. KNN clustering resulted in a total of 8 clusters of EC, **Figure 1B**. EC canonical markers were used to identify the vascular subtype of each cluster. One lymphatic (PROX1, LYVE1, PDPN), two arteriole (EFNB2, HEY1), two capillary (RGCC, KDR), and three venule (ACKR1, SELP) EC clusters were identified, **Figure 1C-D**.

**Figure 1.**
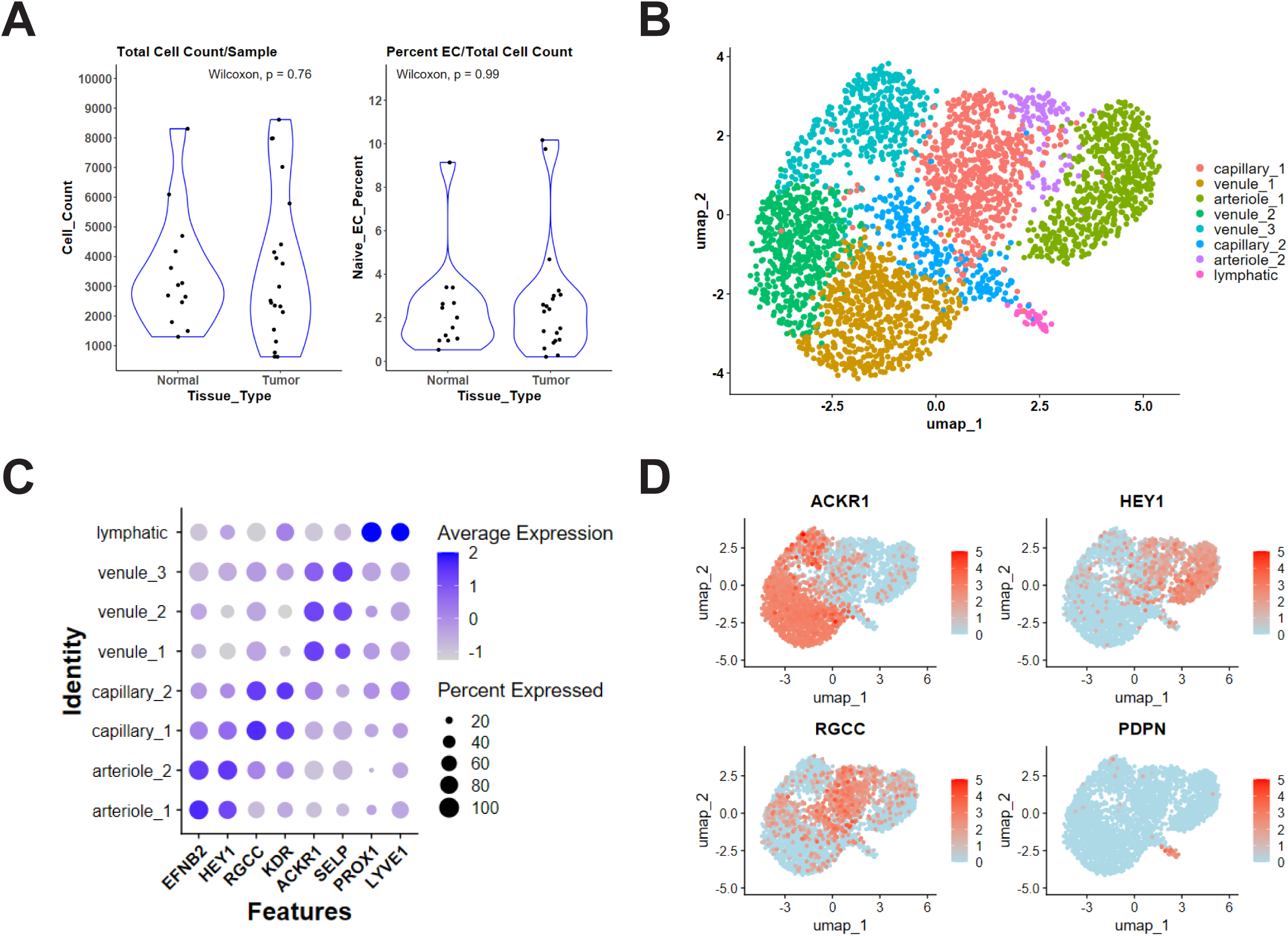
Analysis and clustering of normal and malignant breast tissue endothelial cells. **A.** Comparison of total cell count per patient sample (left) and percentage of endothelial cells of total cells from normal and tumor samples (right). **B.** Clustering of merged and integrated normal tissue endothelial cells (NEC) and tumor tissue endothelial cells (TEC) demonstrating eight clusters. **C-D.** Endothelial subtype determination by canonical gene expression reveals two arteriole, two capillary, three venule, and one lymphatic cluster. **C.** Dot plot comparison of canonical endothelial cell gene expression for arteriole (EFNB2, HEY1), capillary (RGCC, KDR), venule (ACKR1, SELP), and lymphatic (PROX1, LYVE1) subtypes across all clusters. Dot color reflects gene expression level and dot size represents the percentage of cells expressing marker genes within each cluster. **D.** Feature plot comparing canonical endothelial cells gene expression for venule (ACKR1), arteriole (HEY1), capillary (RGCC), and lymphatic (PDPN) subtypes across all cells. The dot color reflects the gene expression level. P-values were determined by the Wilcoxon rank sum test.

### Differential Distribution of Breast Endothelial Cell Subtypes from Normal and Tumor Tissue

Following endothelial cell selection, cell subtype distribution was assessed to determine whether tumor-associated EC exhibited altered vascular phenotypes compared to normal tissue. Analysis of NEC and TEC subtype distribution revealed distinct shifts in vascular composition, **Figure 2A**. NEC exhibited a balanced representation of capillary and venule subtypes, whereas TEC populations were predominantly venule-derived, suggesting a tumor-associated shift toward post-capillary venule enrichment, **Figure 2B**, **Table 2**. In total, the proportional distribution of endothelial cell subtypes was significantly different between normal and tumor breast tissue samples as determined by Chi-squared analysis, p = 0.0246. Individual cluster analysis showed that arteriole_1, venule_1, and venule_2 clusters were significantly increased in TEC compared to NEC, while the venule_3 cluster was significantly increased in NEC representing > 85% of all the normal tissue venule subtype endothelial cells, **Figure 2C**. Venule subtype TEC were > 92% contained in the venule_1 and venule_2 clusters. No difference was observed in the comparison of arteriole_2, capillary_1, capillary_2, or lymphatic subtype clusters between NEC and TEC.

**Figure 2.**
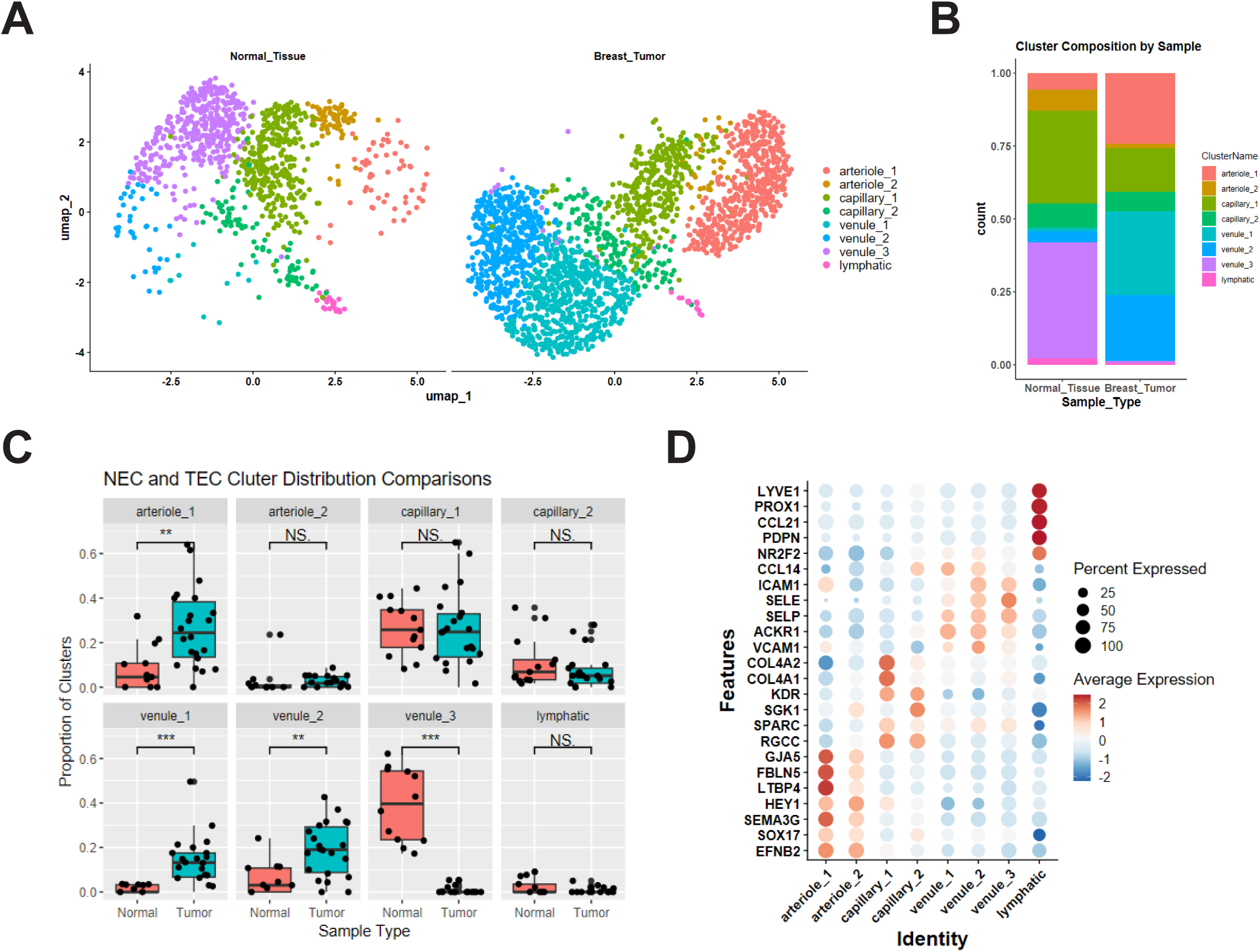
Comparison of cluster endothelial cell distribution by tissue source – normal or malignant breast tissue. **A.** Endothelial cell clusters split by normal or tumor tissue type – normal tissue or breast tumor). **B.** Cluster composition by sample type – normal tissue or breast tumor – for all samples. **C.** Direct comparison of the proportion of cells per cluster from each patient sample across normal (NEC) and tumor (TEC) endothelial cells. P-values were determined by the Wilcoxon rank sum test. Non-significant (ns) = p ≥ 0.05, significant (*) = p ≤ 0.05, (**) p = p ≤ 0.01, (***) p = p ≤ 0.001. **D.** Endothelial cell cluster characterization by gene expression. Dot plot of gene expression of previously defined endothelial cell subtypes across clusters. Dot color reflects gene expression level and dot size represents the percentage of cells expressing genes within each cluster.

**Table 2:** Endothelial cell subtype distribution between normal and tumor tissues.

| **Endothelial Cell Subtype Distribution** |  |  |
| --- | --- | --- |
| **Type** | **Normal (%)** | **Tumor (%)** |
| **Art** | 12.9 | 25.8 |
| **Cap** | 40.3 | 21.8 |
| **Ven** | 44.6 | 52 |
| **Lymph** | 2.2 | 0.6 |
| **Chi-squared analysis** |  |  |
| **TEC vs NEC - P value = 0.0246** |  |  |

A deeper evaluation of cluster-specific gene expression revealed differences in endothelial phenotypes of overlapping parental clusters, **Figure 2D**. Arteriole_2 cluster displayed decreasing levels of arteriole markers, including FBLN5, GJA4, and GJA5, complemented with increasing expression of capillary markers, including RGCC, which is consistent with a post-arterial capillary phenotype compared to arteriole_1. The capillary_1 cluster displayed increased gene expression associated with an angiogenic capillary phenotype, including COL4A1, and COL4A2, compared to capillary_2 cluster cells.[62,63] Capillary_1 also displayed markers of endothelial tip cells, the leading cells of vascular sprouts during angiogenesis, including ESM1, PXDN, APLN, and ANGPTL2.[64,65] Meanwhile, capillary_2 demonstrated increased levels of FABP4, FABP5, and CD36, which is indicative of a fatty acid uptake and lipid metabolism phenotype, as previously described in breast tumor endothelial cells.[39] Venule_1 and venule_2 clusters display markers of activated post-capillary venules; conversely, venule_3 displays venule markers consistent with a non-activated, quiescent post-capillary phenotype. The significant increase in the proportion of venule_1 and venule_2 cluster cells in TEC compared to NEC suggests a shift toward an activated post-capillary venule phenotype.

### Breast TEC gene expression patterns are distinct from NEC

To identify molecular drivers underlying TEC phenotypic shifts, differential gene expression analysis was conducted based on observed TEC subtype differences. NEC and TEC data were grouped by sample type (NEC vs TEC) using a pseudobulk method to aggregate the scRNA-seq data to reduce the risk of false discoveries and effects from confounding variables.[48,51,52,66] A total of 1188 differentially expressed genes (DEGs) were identified (adjusted p-value < 0.05, **Supplemental Table 1**). TECs exhibited many of the up-regulated genes, with 1058 DEGs up-regulated compared to only 130 up-regulated DEGs in NEC. Among the top up-regulated DEGs in TEC were ARHGDIB, RAMP2, JAM2, IGF2, PTP4A3, ALDOA, and CD34, indicating activation of pathways related to angiogenic remodeling, extracellular matrix (ECM) reorganization, and endothelial permeability.[39,67–72] In contrast, MALAT1, MT2A, TM4SF1, FTL, CCL2, CXCL8, CXCL1, and STC1 were preferentially expressed in NEC or down-regulated in TEC, suggestive of a loss of immune surveillance and altered vascular integrity within tumor-associated vasculature. **Figure 3A**.[17,21,73–79] Gene ontology (GO) enrichment analysis of the top 500 TEC up-regulated DEGs identified several angiogenic mechanisms, extracellular matrix (ECM) organization, endopeptidase regulation, complement activation, and arteriole development as enhanced biological processes in TEC compared to NEC, **Figure 3B**. Further, a pre-ranked gene list from NEC/TEC differential expression analysis was used in a Gene Set Enrichment Analysis (GSEA) to determine which MSigDB hallmark gene sets were enriched, **Figure 3C**. Genes that are upregulated in response to interferon gamma and interferon alpha were enriched in TEC while genes associated with inflammatory response and genes regulated by NF-kB in response to TNF were enriched in NEC compared to TEC, **Figure 3D**. To evaluate specific transcriptomic modulation, a list of 200 NF-kB target genes regulated by TNFA signaling from MSigDB [55] was assessed for differential expression between NEC and TEC, **Figure 3E**. Several NF-kB target genes were observed to be repressed in TEC compared to NEC, including IL6, CCL2, and NFKBIA, further supporting the presence of aberrant NF-kB signaling in TEC. Inflammatory signaling molecules were repressed in TEC when compared to NEC, as well as SELE, a key adhesion molecule that enables inflammatory cell infiltration into the TME. TNFSF10 was observed to be increased in TEC compared to NEC, which was previously shown to promote endothelial cell-mediated suppression of CD8+ T-cells in breast cancer.[80]

**Figure 3.**
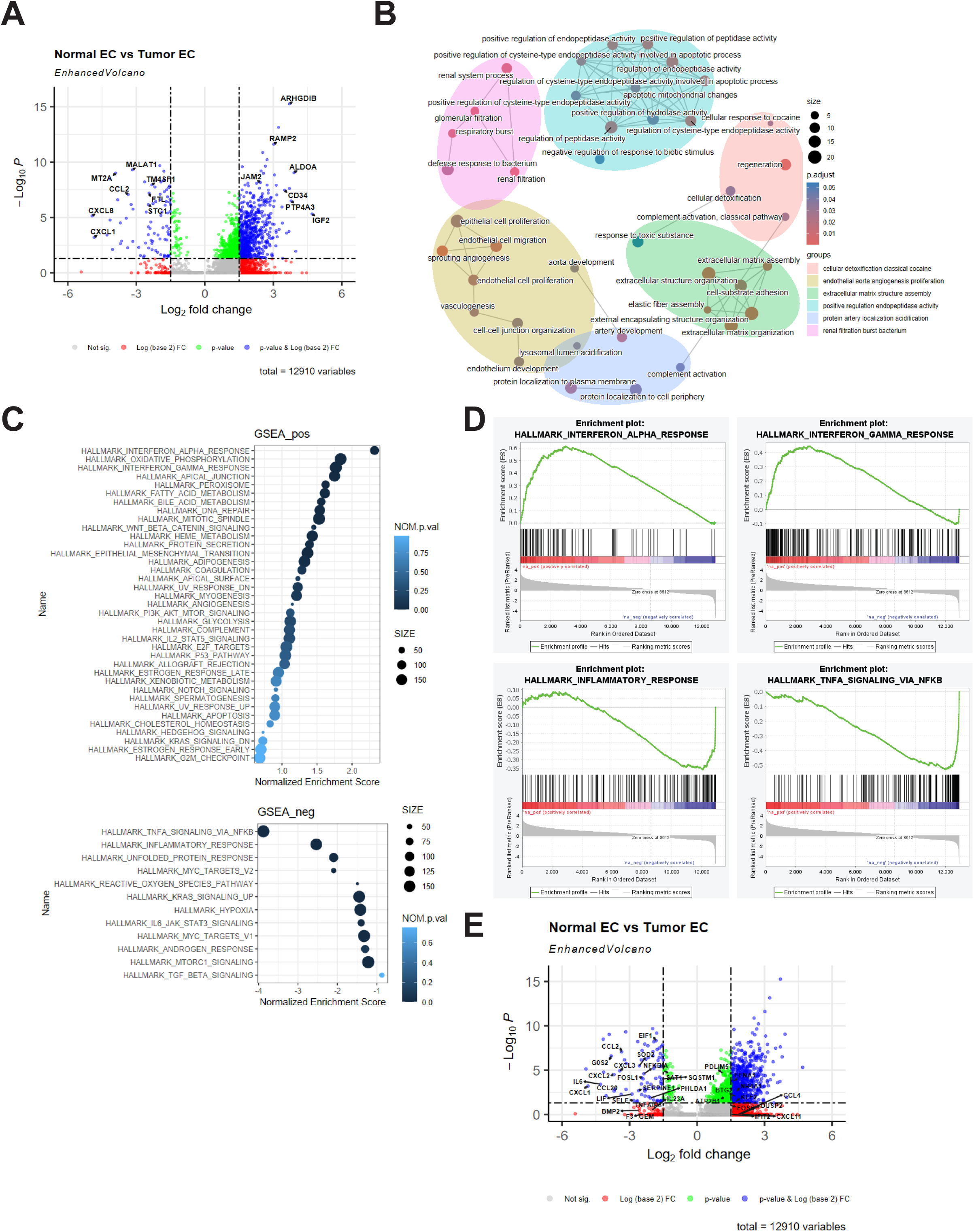
Differential gene expression between NEC and TEC. **A.** Volcano plot showing differentially expressed genes by Log2 fold change and –Log10 p-value. **B.** Enrichment network map of the top 50 categories from gene ontology analysis of the top 500 TEC up-regulated genes. Six identified clusters of biological processes are grouped by color. Size of dot represents the number of genes per category and color represents the adjusted p-value. **C.** Gene Signature Enrichment Analysis (GSEA) of a pre-ranked list of genes from the differential gene expression compared to the MSigDB Hallmark gene sets. Positively enriched (top panel) and negatively enriched (botton panel). Size of dot represents the number of genes per category and color represents the nominal p-value. **D.** Enrichment plots for four hallmark gene sets from the GSEA. **E.** Volcano plot of differential gene expression analysis of NEC and TEC with representative NFKB genes regulated by TNFA labeled.

### Identification of Tumor Endothelial Markers in Breast Tumors

The expression of established endothelial cell markers was evaluated in the pseudobulk analysis to validate the approach, **Figure 4A**. Endothelial markers including PECAM1, CDH5, CD34, and VWF, were increased in TEC compared to NEC. F8, ACE, and VCAM1 were expressed at low levels in both NEC and TEC though all were significantly increased in TEC. CLDN5 was strongly expressed in both NEC and TEC and not observed to change between the two. Finally, SELE and ICAM1 displayed decreased expression in TEC compared to NEC. The data from breast NEC and TEC was further evaluated for the expression of tumor endothelial markers (TEM) previously identified in colorectal cancer and lung cancer.[81,82] There was little to no expression of the majority of these TEMs detected, and they were not identified as upregulated in the pseudobulk DEG analysis, **Supplemental Table 1**. While ADGRA2, PLXDC1, CHN1, MXRA5, LCP2, and POSTN were significantly increased in TEC compared to NEC, the overall expression was minimal, **Supplemental Figure 1.** Further, several genes, including HSPA6, RPS11, STC1, TNFAIP6, and PMAIP1 were significantly downregulated in TEC compared to NEC. This data supports the known heterogeneity of tumor endothelial cells between cancer subtypes, extending this observation to TEMs.

**Figure 4.**
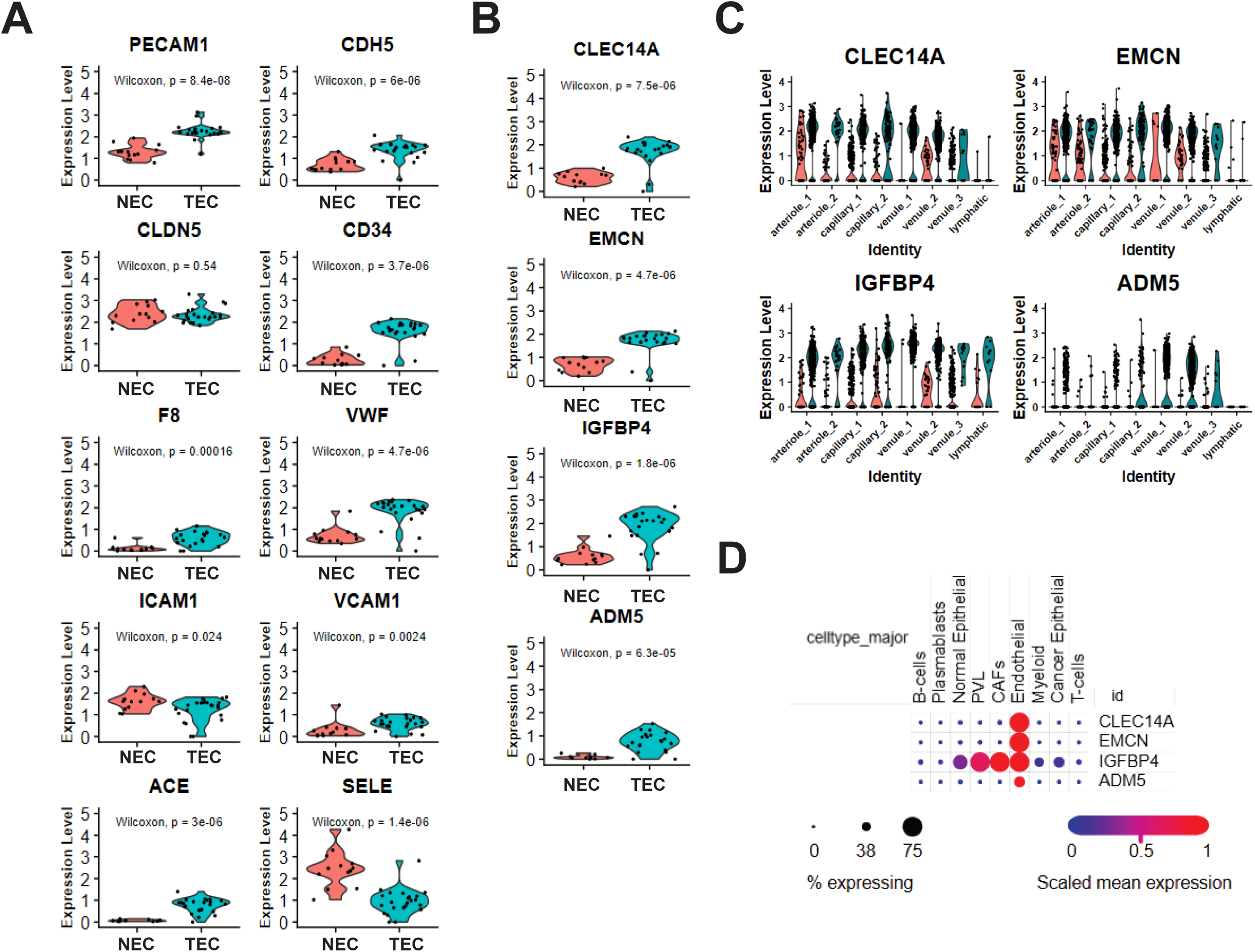
Expression of endothelial genes in NEC and TEC. **A.** Expression of classical endothelial marker genes. **B.** Expression of identified breast TEC specific genes, CLEC14A, EMCN, IGFBP4, and ADM5 in NEC and TEC. **C.** Violin plots of NEC and TEC expression of CLEC14A, EMCN, IGFBP4, ADM5 across each endothelial cell cluster, split by sample type (NEC (red), TEC (blue)). **D.** Expression of CLEC14A, EMCN, IGFBP4, ADM5 in the initial Wu et al major cell type clusters. Dot color reflects gene expression level and dot size represents the percentage of cells expressing genes within each cluster. P-values were determined by the Wilcoxon rank sum test.

To evaluate breast TEC for specific TEMs, representative genes identified by DEG were evaluated for TEC expression and distribution. CLEC14A, EMCN, IGFBP4, and ADM5 were significantly increased in TEC compared to NEC, **Figure 4B**. CLEC14A, EMCN, and IGFBP4 were found to be expressed at low frequency in NEC, while ADM5 was not expressed in NEC. Additionally, CLEC14A, EMCN, and IGFBP4 were expressed in all clusters except for the lymphatic cells, displaying the highest expression in tumor venule clusters. ADM5 was selectively expressed by the two TEC majority venule clusters, **Figure 4C**. To confirm the specificity of the candidate TEM genes, the expression of CLEC14A, EMCN, IGFBP4, and ADM5 was evaluated in the original analysis from Wu et al. in the Broad Institute Single Cell Portal.[46] Expression of CLEC14A, EMCN, and ADM5 was specific to endothelial cells, **Figure 4D**. IGFBP4 was expressed in endothelial cells as well as perivascular-like cells, cancer associated fibroblasts, and both normal and malignant epithelial cells. To extend the cell type specificity analysis across multiple cancer types, pan-cancer scRNA-seq data was assessed in the Curated Cancer Cell Atlas (3CA), **Supplemental Figure 2**. Expression of CLEC14A, EMCN, and ADM5 remained highly specific to endothelial cells across many cancer types, with expression of ADM5 present in a minor percentage of the endothelial cell population. IGFBP4 was expressed in endothelial cells as well as pericytes, fibroblasts, and some epithelial and malignant cells. Finally, to expand the analysis of TEC upregulated genes, expression levels of CLEC14A, EMCN, IGFBP4, and ADM2 were compared to all other genes in the METABRIC and TCGA breast cancer bulk RNA-seq datasets and Spearman’s correlation was determined, **Supplemental Table 2**. CLEC14A and EMCN were strongly correlated with multiple other endothelial cell associated genes, including CDH5, VWF, and CD34. IGFBP4 and ADM5 were only moderately correlated with endothelial cell-associated genes. IGFBP4 also correlated with ECM-related genes, LAMB2 and LAMA2, while ADM5 was correlated with specifically the venule-type endothelial cell genes ACKR1 and CCL14. Taken together, the RNA expression of CLEC14A, EMCN, and ADM5 is highly specific to TEC across cancer types and are candidate TEM, while IGFBP4 is not specific to TEC.

### Increased ADM5 Expression is Associated with Poor Patient Survival

To assess the potential translational value of identified TEC up-regulated genes, survival analysis based on gene expression was performed by univariate Kaplan–Meier regression analysis. A total of 2,976 breast cancer patients were assessed for overall survival (OS) compared to CLEC14A, EMCN, IGFBP4, and ADM5 gene expression. No significant effect of high vs low expression levels was observed in the overall BC cohort, **Supplemental Figure 3**. Conversely, when the basal-only PAM50 subtype patients (n=309) were evaluated for survival based on gene expression, high expression of ADM5 was associated with a significantly reduced OS (HR = 2.27, 95% CI 1.37-3.75; p = 0.0001), **Figure 5A**. No significant change in OS was observed for CLEC14A, EMCN, or IGFBP4. This data was confirmed in the TCGA Breast Cancer dataset of basal subtype in cBioPortal (HR = 2.57, 95% CI 1.11-5.92, p = 0.0395), where high ADM5 expression was associated with reduced OS, **Supplemental Figure 4**. Since ADM5 was observed to be highly specific to the endothelial compartment across multiple tumor types, additional cancer types were evaluated for impact on OS. In addition to basal-type breast cancer patients, patients with gastrointestinal-related cancers including gastric, stomach, and colon cancer as well as renal clear cell carcinoma and ovarian cancer all were found to have significantly decreased OS with high ADM5 expression, **Figure 5B**. The association of ADM5 expression with poor clinical outcomes suggests a potential role in tumor immune evasion or therapeutic resistance mechanisms.

**Figure 5.**
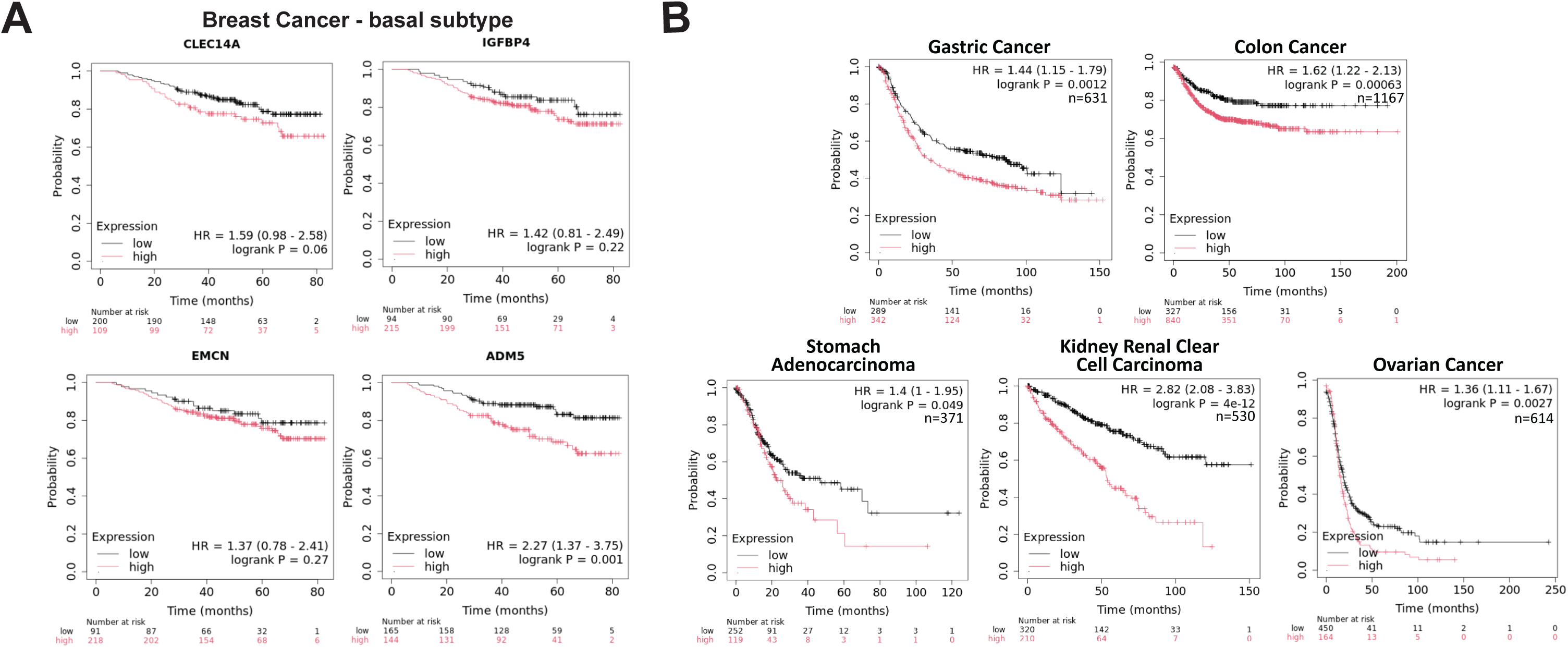
**A**. Correlation of CLEC14A, EMCN, IGFBP4, and ADM5 gene expression to overall survival in breast cancer patients with the PAM50 basal subtype (n=309). **B.** Correlation of ADM5 gene expression to overall survival in patients with gastric cancer (n=631), colon cancer (n=1167), stomach adenocarcinoma (n=371), kidney renal clear cell carcinoma (n=530), and ovarian cancer (n=614). Cox proportional hazards regression analysis was used to calculate hazard ratio (HR) h with 95% confidence intervals and log-rank p-value.

To further develop the context of ADM5 expression in TEC, ADM5-associated signaling pathway genes were evaluated for expression in the breast NEC and TEC dataset, **Figure 6A, Supplemental Figure 5**. ADM and ADM5 were expressed across multiple clusters of endothelial cells, no other pathway-related ligand genes were observed including ADM2, CALCA, CALCB, and IAPP. The receptor-associated proteins, RAMP2 and RAMP3 were expressed across all TEC subtype clusters, while RAMP1 was not detected. CALCR was not expressed, CRCP was minimally expressed, and CALCRL was widely expressed in normal and tumor samples across all EC clusters. Further, ADM5 (2.9 log2FC), RAMP2 (3.0 log2FC), and RAMP3 (3.1 log2FC) were significantly increased in TEC compared to NEC in differential gene expression analysis, **Supplemental Table 1**. CALCRL was expressed at similar levels between NEC and TEC and broadly expressed across endothelial subtypes. Finally, when evaluated for translational impact, similar to ADM5, high RAMP2 gene expression was found to be associated with significantly reduced basal breast cancer patient OS, **Figure 6B**. High ADM expression was associated with improved OS, while RAMP3 and CALCRL expression showed no significant correlation to OS. Cell type specifity was evaluated in the original Wu et al. analysis in the Broad Institute Single Cell Portal. ADM displayed broad cell type expression while CALCRL, RAMP2, and RAMP3 were highly specific to endothelial cells, **Supplement Figure 5**. The specificity of cancer cell type expression of ADM, CALCRL, RAMP2, and RAMP3 was also assessed in the 3CA dataset and revealed a similar pattern of expression across cancer types (data not shown). Finally, ADM5 gene expression was evaluated for correlation to immunotherapy outcomes in KM Plotter.[83] Interestingly, ADM5 high expression was associated with significantly reduced PFS and OS in PD1 and CTLA4-treated patients across multiple cancer types. Conversely, PDL1-treated patients displayed a significantly increased PFS and OS, **Figure 6C**.

**Figure 6.**
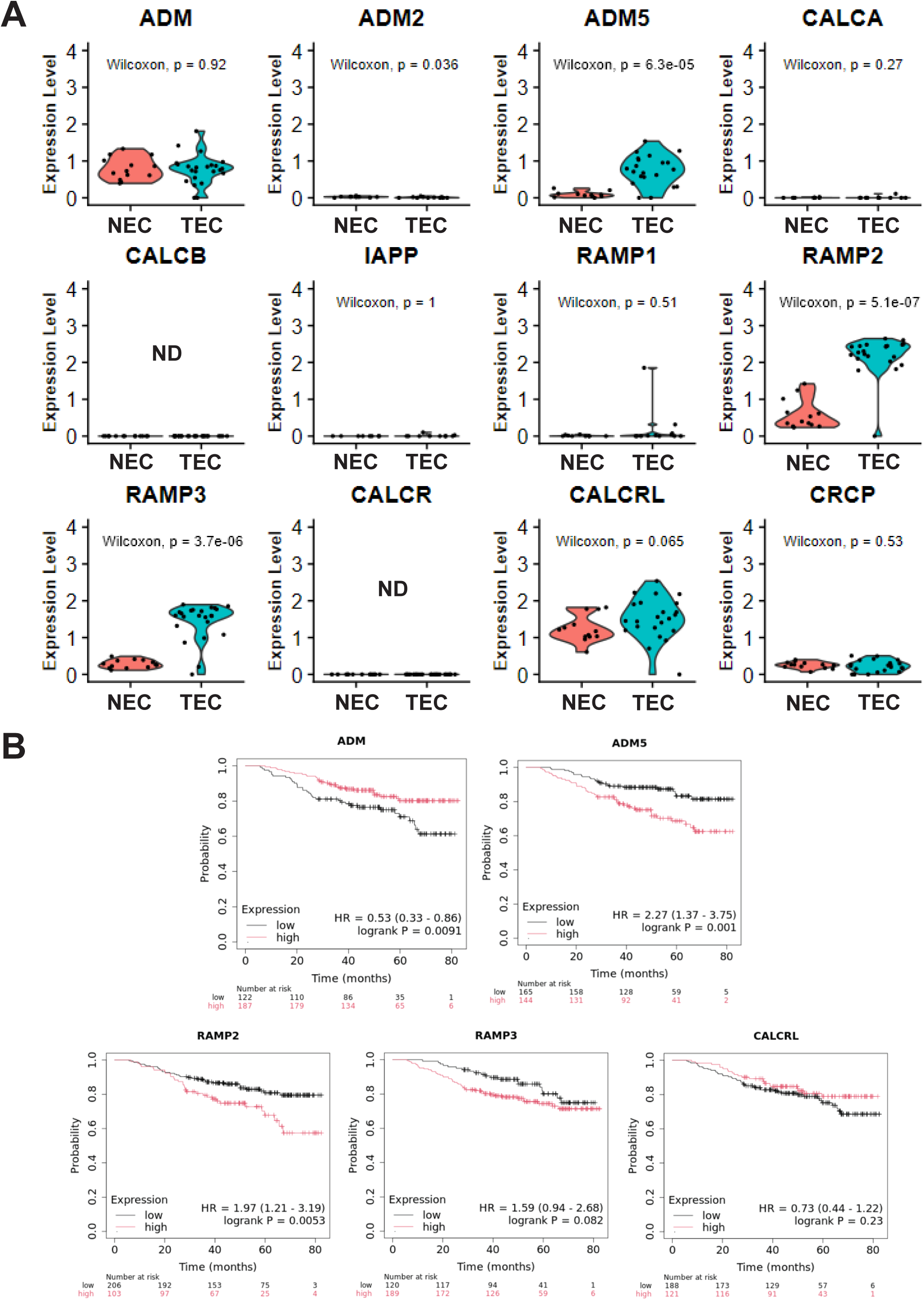
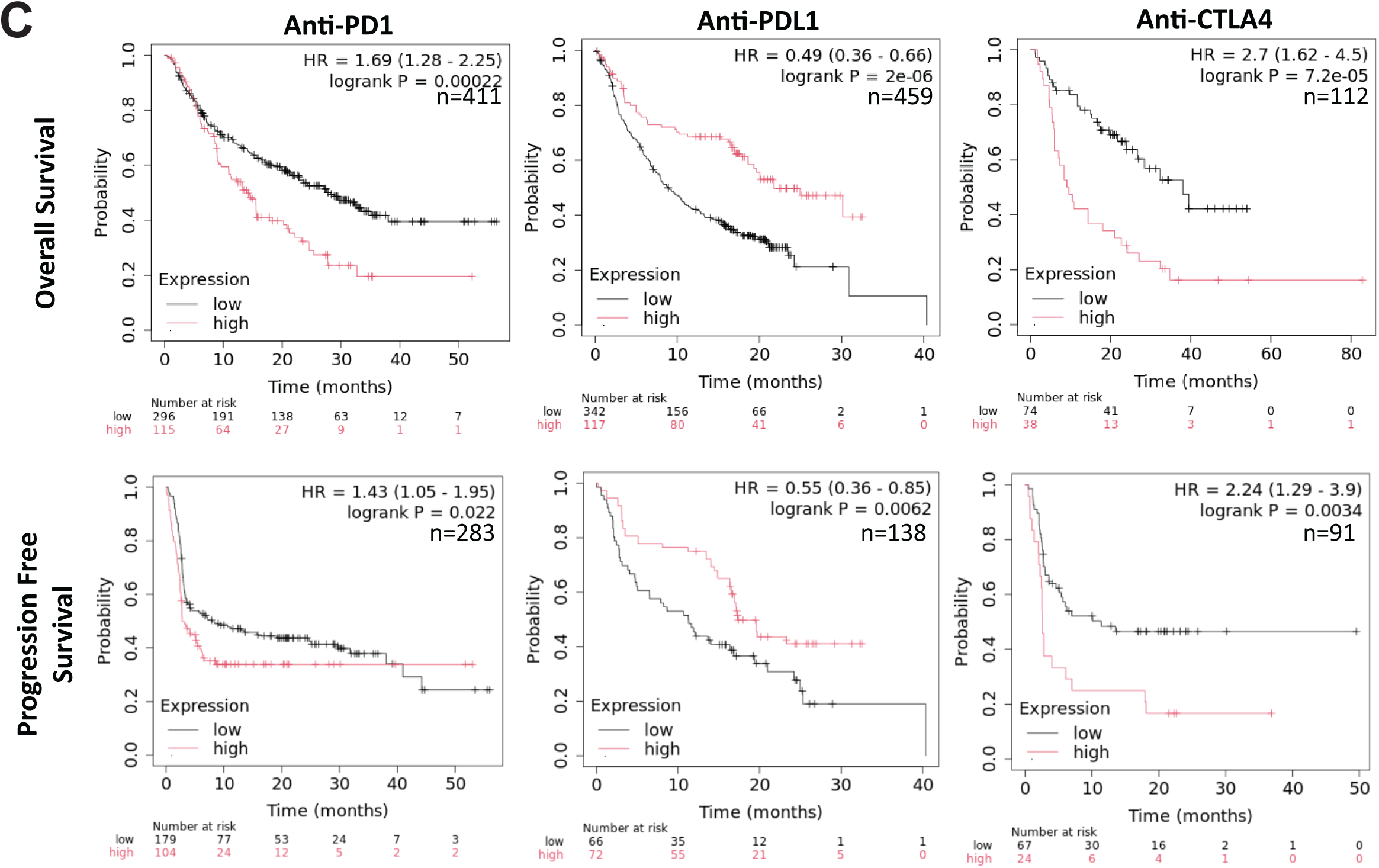
Gene expression and cancer patient survival correlation analyses of ADM5 family genes. **A.** Expression of ADM, ADM2, ADM5, CALCA, CALCB, IAPP, RAMP1, RAMP2, RAMP3, CALCR, CALCRL, CRCP in NEC and TEC. **B.** Correlation of ADM, ADM5, RAMP2, RAMP3, and CALCRL gene expression to overall survival in breast cancer patients with the PAM50 basal subtype (n=309). **C.** Correlation of ADM5 gene expression in anti-PD1, anti-PDL1, and anti-CTLA4 to overall survival and progression-free survival in cancer patients. P-values were determined by the Wilcoxon rank sum test. Cox proportional hazards regression analysis was used to calculate hazard ratio (HR) h with 95% confidence intervals and log-rank p-value. ND = not detected.

## Discussion

This work established a unique scRNA-seq EC atlas of normal and tumor-derived breast endothelial cells providing a unique dataset to compare NEC and TEC populations. Rather than use unsupervised clustering of total tissue cell populations or experimental enrichment of endothelial cells via magnetic or FACS-based methods, this study utilizes canonical gene expression to purify the cells of interest. An evaluation of the endothelial cells identified via unsupervised clusters from the original analysis of GSE176078 revealed that cells in the endothelial cluster also expressed markers of smooth muscle cells, pericytes, fibroblasts, and immune cells (data not shown). The method applied in this study provides a more stringent evaluation of endothelial cells from breast tissue including normal and tumor derived, treatment naïve samples. It should be noted that the exclusion of cells expressing ACTA2 and COL1A1 would also remove any endothelial cells undergoing endothelial-to-mesenchymal transition (EndoMT) from this dataset. EndoMT results in decreased expression of classical endothelial markers and induced expression of mesenchymal markers leading to dramatic phenotypic and functional changes in endothelial cells.[84] An investigation of endoMT would therefore require an alternative method of cell identification not utilized in this work.

A comparison of endothelial cell subtypes revealed that TEC consists of a majority venule subtype and displays a significant difference in the distribution of arteriole, capillary, and venule EC compared to NEC subtypes. While many differences in TEC phenotypes have been described, no observations have been made on the distribution of EC subtypes in normal and tumor breast tissue. The reduction in capillary subtype cells observed in TEC is consistent with observations reported in normal lung and lung tumor endothelial cell analyses.[85] The venule profiles of NEC and TEC were strikingly different. The venule_1 and venule_2 clusters were comprised of nearly all TEC-derived, while the venule_3 cluster was mostly NEC-derived. This suggests that while TEC are generally different from NEC, the venule population may be a major contributor to these differences, especially given the critical functional role of venule-type cells in immune cell infiltration into tumors. NEC displayed expression of immune cell adhesion markers SELE and ICAM1, which were decreased in the TEC population consistent with the reported altered phenotype of TEC and reduction of molecules required for trans-endothelial migration of immune cells. This reduction in immune cell recruitment and expression of adhesion markers on TEC is in alignment with an endothelial anergy phenotype where chronic stimulation results in the non-responsiveness of the tumor vasculature to inflammatory stimulation and downregulation of adhesion molecules.[17,28,77,86–88] The data in this study revealed several additional genes repressed in TEC compared to NEC, including IL6, CXCL1, CXCL8, and CCL2. The reduction in expression of these proinflammatory signals in TEC can directly limit the recruitment of neutrophils, monocytes, macrophages, and other immune cells to the TME.[89,90] Continued exploration of breast TEC-diminished genes will be required to determine the specific contribution of these genes to mechanisms of endothelial anergy in breast cancer. Extension of this analysis to include cell – cell interaction and signaling that may drive EC anergy and direct tumor immune status.

Several of the TEC suppressed genes; SELE, IL6, CXCL1, CXCL8, CCL2, and CCL20 are downstream targets of NF-κB.[91] The observation that these genes are downregulated in breast TEC compared to NEC indicates that breast tumor vasculature displays aberrant NF-κB signaling. NFKBIA, a negative regulator of NF-κB, was also downregulated in TEC vs NEC which suggests that the suppression of NF-κB in TEC is potentially independent of canonical NF-κB pathway regulation and likely due to alternative compensatory mechanisms. NF-κB is an angiostatic regulator known to contribute to the maintenance of vascular homeostasis and endothelial integrity.[92,93] In fact, in a study on angiostatic agents, NF-κB inhibition prevented the reversal of endothelial anergy in a leukocyte adhesion assay.[93] It is possible that breast TEC anergy may be driven, at least partially, by aberrant NF-κB signaling. Targeting NF-κB signaling specifically in TEC should be investigated as a method to reactivate tumor vasculature and improve patient outcomes to cancer therapies, particularly immunotherapy regimens.

Identification of TEC-specific markers can lead to the identification of biomarkers or novel therapeutic targets. The expression of CLEC14, EMCN, IGFBP4, and ADM5 were found to be significantly increased in breast TEC compared to NEC. CLEC14 has been previously identified as a TEM in various solid tumors, including breast.[94,95] CLEC14A, a C-type lectin-like type I transmembrane protein, has been shown to contribute to tumor angiogenesis, and targeting CLEC14a with CAR-T or antibody therapeutics resulted in reduced tumor growth in preclinical models of pancreatic, lung, and hepatic cancer.[96,97] The identification of CLEC14a expression in breast TEC confirms the robustness of our informatics approach and supports the validity of new and less described markers identified here. EMCN, endomucin or MUC14, is a transmembrane *O*-sialylated protein known to be expressed by endothelial cells with enrichment in the capillary and venous subtypes. It has been shown to contribute to pathological angiogenesis and the regulation of cell adhesion and extracellular matrix interactions of endothelial cells.[98–102] The application of EMCN as a TEM has not been described. In this dataset, EMCN was highly specific to TEC compared to NEC and was observed to be endothelial cell-specific in pan-cancer cell type analysis. Overexpression of EMCN has been reported to decrease immune cell infiltration under inflammatory conditions by reducing endothelial cell-leukocyte interactions, and EMCN KO resulted in increased immune cell infiltration.[103,104] Taken together, these data suggest that increased EMCN on tumor vasculature may be an adaptation to enable tumor progression and evade immune response. Blocking or degrading EMCN on TEC may offer a therapeutic opportunity to improve immune cell infiltration into solid tumors. IGFBP4 is a secreted protein that can bind to and inhibit IGF activity while also displaying IGF-independent functions including roles in bone growth and angiogenesis.[105] While IGFBP4 was strongly up-regulated in TEC compared to NEC, expression was not specific to the endothelial compartment with observed expression in pericytes, fibroblasts, and tumor cells across multiple cancer cell types. Given this data, IGFBP4 does not appear to be a specific TEM candidate, however further investigation may be warranted to discern the role of IGFBP4 in the TME given that previous data has shown that inhibition of IGFBP4 cleavage, and therefore reduction in local IGF signaling, reduced tumor growth and increased endothelial cell apoptosis in a mouse model of breast cancer.[106] ADM5 represents a novel TEM candidate that has not previously been reported in TECs. This analysis revealed ADM5 to be highly specific to the breast tumor endothelial cell population, with pronounced enrichment in the venule endothelial subtype. Importantly, ADM5 expression was restricted to endothelial cells within the source tumor tissue and pan-cancer transcriptomic profiling confirmed its absence in all other cell types. This specificity underscores ADM5’s potential as a TEC-selective marker with translational relevance. High expression of ADM5 was correlated to decreased OS in basal-type breast cancer patients and other tumor types including gastric, renal, and ovarian cancer. Notably, basal-type breast tumors are among the most angiogenic subtypes, alongside inflammatory breast tumors, exhibiting heightened vascular proliferation rates relative to other breast cancer subtypes.[107,108] This distinct vascular phenotype may underlie the observed correlation between elevated ADM5 levels and poor prognosis in basal-type breast cancer patients. Given that other adrenomedullin ligands contribute to angiogenic signaling and endothelial remodeling, ADM5 may similarly influence tumor vascular dynamics, potentially exacerbating disease progression.[109,110]

ADM5 is a poorly characterized member of the calcitonin gene-related peptide (CGRP) family that includes CGRP, calcitonin (CT), amylin (AMY), and adrenomedullin (ADM). The adrenomedullin family consists of ADM1, ADM2, and ADM5. ADM1 and AMD2/IMD have direct effects on endothelial cell biology and have been shown to signal through calcitonin receptor-like receptor (CALCRL) in complex with one of three receptor activity-modifying proteins (RAMP), which alter the ligand affinity and signaling specificity.[111,112] RAMP2 participates in vascular endothelial signaling while RAMP3 directs lymphovascular effects.[67,113] Expression data supports the hypothesis that ADM5 may signal in an autocrine fashion through CALCRL complexed with RAMP2 or RAMP3, as these displayed similar cell-type specificity. The observation that only ADM5 and RAMP2 were associated with reduced overall survival in basal-type breast cancer patients suggests that RAMP2 is more likely to contribute to ADM5 signaling in breast cancer. Finally, high ADM5 expression was associated with reduced PFS and OS in patients treated with PD1 and CTLA4 blocking therapies but not with anti-PDL1 treatment, suggesting that ADM5 expression may contribute to reduced efficacy of T-cell mediated checkpoint inhibitor therapeutics while not impeding cancer cell targeted therapeutics. PD1, PDL1, and CTLA-4 displayed little to no expression in NEC and TEC (data not shown) suggesting that the change in OS is not a direct impact on TEC therapeutic targeting. A recent study demonstrated that the related and signaling peptide, ADM, contributed to the inhibition of therapeutic response in anti-PD1 immunotherapy combined with VEGFR2 inhibition in biliary tract cancer patients through direct effects on endothelial cell function.[114] Interestingly, ADM has been shown to interface with the NFKB pathway in rat retinal endothelial cells where ADM treatment inhibited VEGF-induced inflammatory responses via inhibition of NFKB signaling.[109] Further, ADM treatment suppressed expression of cell adhesion molecules, including SELE, in lymphatic EC.[115] Taken together, these data highlight the potential importance of similar signaling by ADM5 in breast cancer, suggesting that increased endothelial ADM5 may have a role of immune evasion supported by tumor vasculature. The exact role of ADM5 in TEC biology will need to be further defined with additional studies.

## Limitations of the Study

Potential limitations of this study include common drawbacks of transcriptomic methods, such as tissue processing effects on cell status and quality. It is possible that tissue processing conditions or variability between sample processing methods could impact EC recovery efficiency or directly activate or inhibit pathways in endothelial cells. Additionally, normal and tumor tissue samples are not patient-matched, so some observations may be influenced by patient-specific profiles. Therefore, confirming these results in additional patient cohorts is critical to validate the findings. Further analysis with additional patient samples and expanded cluster analysis could also reveal additional EC subtypes that were not detected in this analysis due to low cell counts. It should also be noted that collection methods of normal samples tend to source tissue from younger women. In fact, the average age of the patient samples from the breast tumor cohort was 56.9y (+/-13.5) while the normal tissue sample patients had an average age of 40.1y (+/-14.8) (p=0.002). Future analyses should aim to collect age-matched tissue samples to reduce potential confounding variables including age-related changes and hormonal effects on endothelial cells

This study evaluated differences between endothelial cells in normal and treatment-naïve malignant breast tissues, providing a snapshot of breast TEC biology prior to the application of therapeutic interventions. Analyzing the effect of therapeutic cancer treatments on TEC transcriptomic profiles is a potential next step in this line of investigation. As with other cell types in the TME, endothelial cells are likely to change and adapt in response to patient cancer treatments. It is imperative to understand how identified TEMs are affected, especially in the context of disease progression and survival outcomes. As patient PFS and OS data are not available for large numbers of treatment-naïve patients, this may confound the survival analyses across cancer types or subtypes.

To substantiate ADM5 as a novel TEM, further validation is required. Specifically, in situ studies are needed to confirm ADM5 mRNA and protein expression, delineate spatial localization within the tumor vasculature, and establish cell type specificity. These investigations will be critical to affirm ADM5’s utility as a TEC-selective marker with translational relevance.

## Summary

Tumor vasculature is a key player in the progression and spread of breast cancer. Chronic inflammation within the TME fosters endothelial anergy, diminishing therapeutic responses. Identifying TEC-specific markers can pave the way for novel anti-angiogenic therapies and deepen our understanding of breast cancer-specific endothelial anergy mechanisms. By rigorously analyzing endothelial cell profiles in normal and malignant breast tissues, this study uncovered significant differences in endothelial subtypes, phenotypic profiles, and highlighted numerous candidate genes for further investigation. Notably, ADM5 was identified as a novel TEC-specific marker with potential implications in tumor immune evasion and therapeutic resistance. Its association with poor patient survival and reduced efficacy of immune checkpoint therapies highlights its relevance in vascular-targeted interventions. Future investigations should explore ADM5’s mechanistic role in TEC function, with an emphasis on therapeutic strategies that reprogram endothelial immune responses.

## Availability of Data and Materials

The datasets analyzed during the current study are available in Gene Expression Omnibus (GEO), GSE176078 and GSE161529.

## Acknowledgements

The authors would like to acknowledge the Computational Biology Core, Institute for Systems Genomics, University of Connecticut for supporting this work.

## List of abbreviations

3CA: Curated Cancer Cell Atlas
ADM: adrenomedullin
AMY: amylin
AAT: anti-angiogenic therapy
BC: breast cancer
CT: calcitonin
CGRP: calcitonin gene-related peptide
DEG: differential expressed gene
EC: endothelial cell
EndoMT: endothelial-to-mesenchymal transition
ECM: extracellular matrix
GSEA: Gene Set Enrichment Analysis
KNN: K-means nearest neighbor
GEO: NCBI Gene Expression Omnibus
NEC: normal endothelial cell
OS: overall survival
pCR: pathological complete response
PAM50: prediction analysis of microarray 50
PFS: progression free survival
RAMP: receptor activity-modifying proteins
scRNA-seq: single cell RNA-seq
TEC: tumor endothelial cell
TEM: tumor endothelial marker
TME: tumor microenvironment
VEGF: vascular endothelial growth factor

## Ethics approval and consent to participate

Not applicable

## Consent for publication

Not applicable

## Competing interests

The authors declare that they have no competing interests

## Funding

This work was funded by UConn Research Enhancement Program and UConn Health Department of Cell Biology/Center for Vascular Biology

## Authors’ contributions

KNP conceptualized the study, conducted data analysis, and wrote the initial manuscript draft. VS supported scRNA-seq processing and contributed to bioinformatics troubleshooting. PAM assisted with statistical validation and provided technical oversight in vascular biology and bioinformatic methodologies. KPC supervised the research, secured funding, and provided critical revisions to the manuscript. All authors reviewed and approved the final version.

## Figure Legends

**Supplemental Figure 1.**
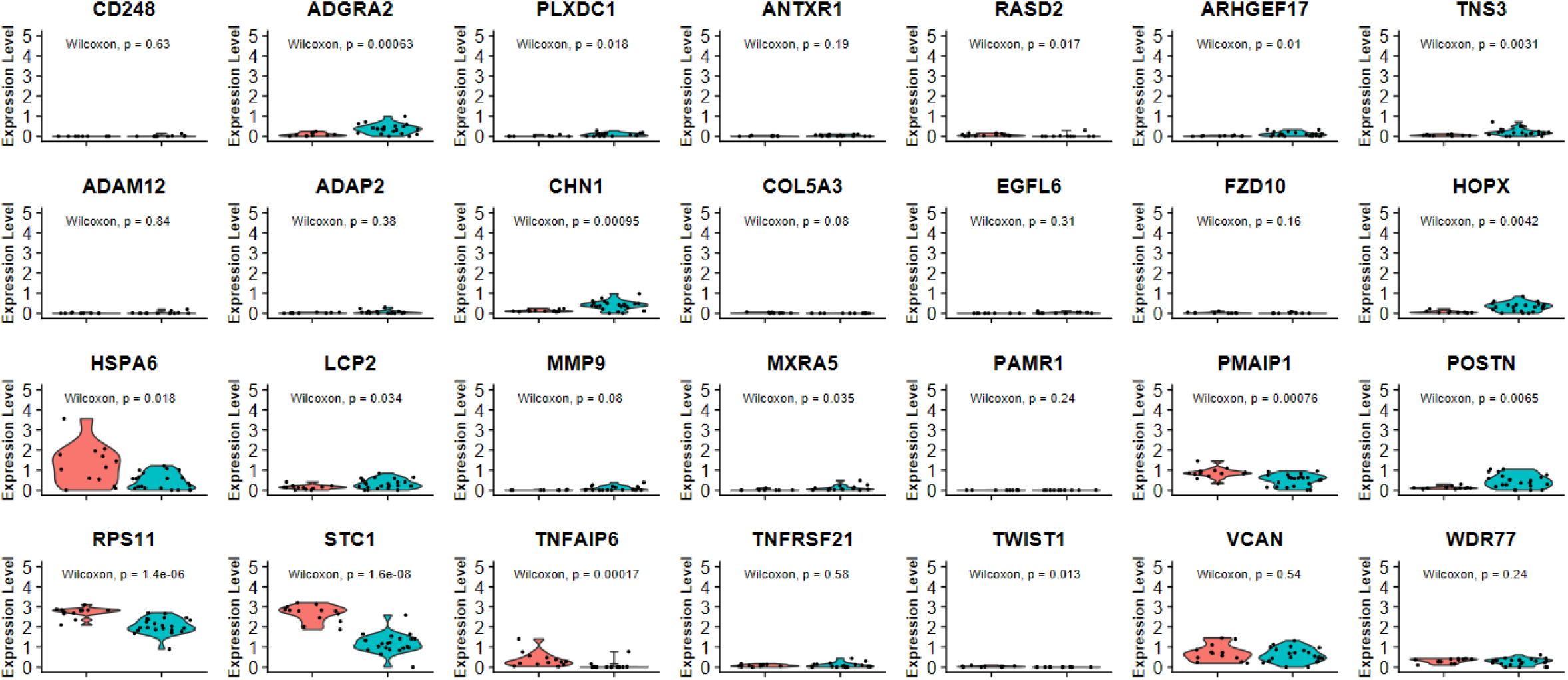
Expression of the previously identified lung cancer and colon cancer TEMs in breast NEC and TEC. P-values were determined by the Wilcoxon rank sum test.

**Supplemental Figure 2.**
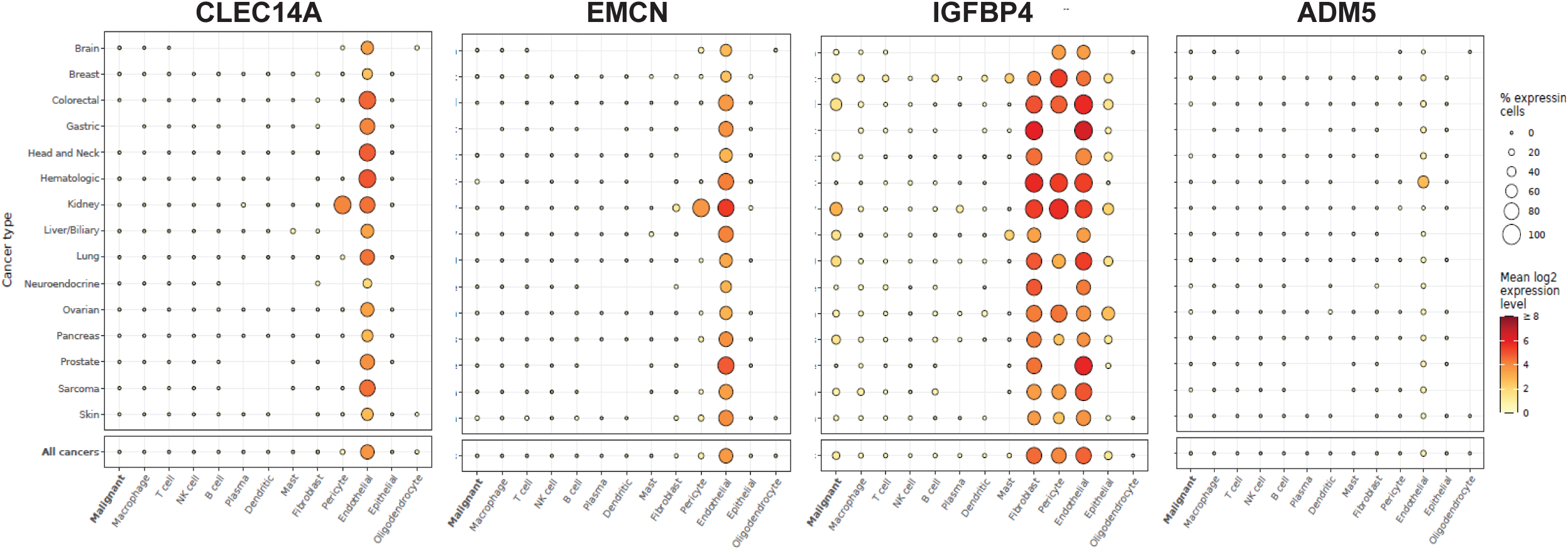
Cell type-specific expression of CLEC14A, EMCN, IGFBP4, and ADM5 in the 3CA pan-cancer single-cell dataset. Dot color reflects gene expression level and dot size represents the percentage of cells expressing genes within each cluster.

**Supplemental Figure 3.**
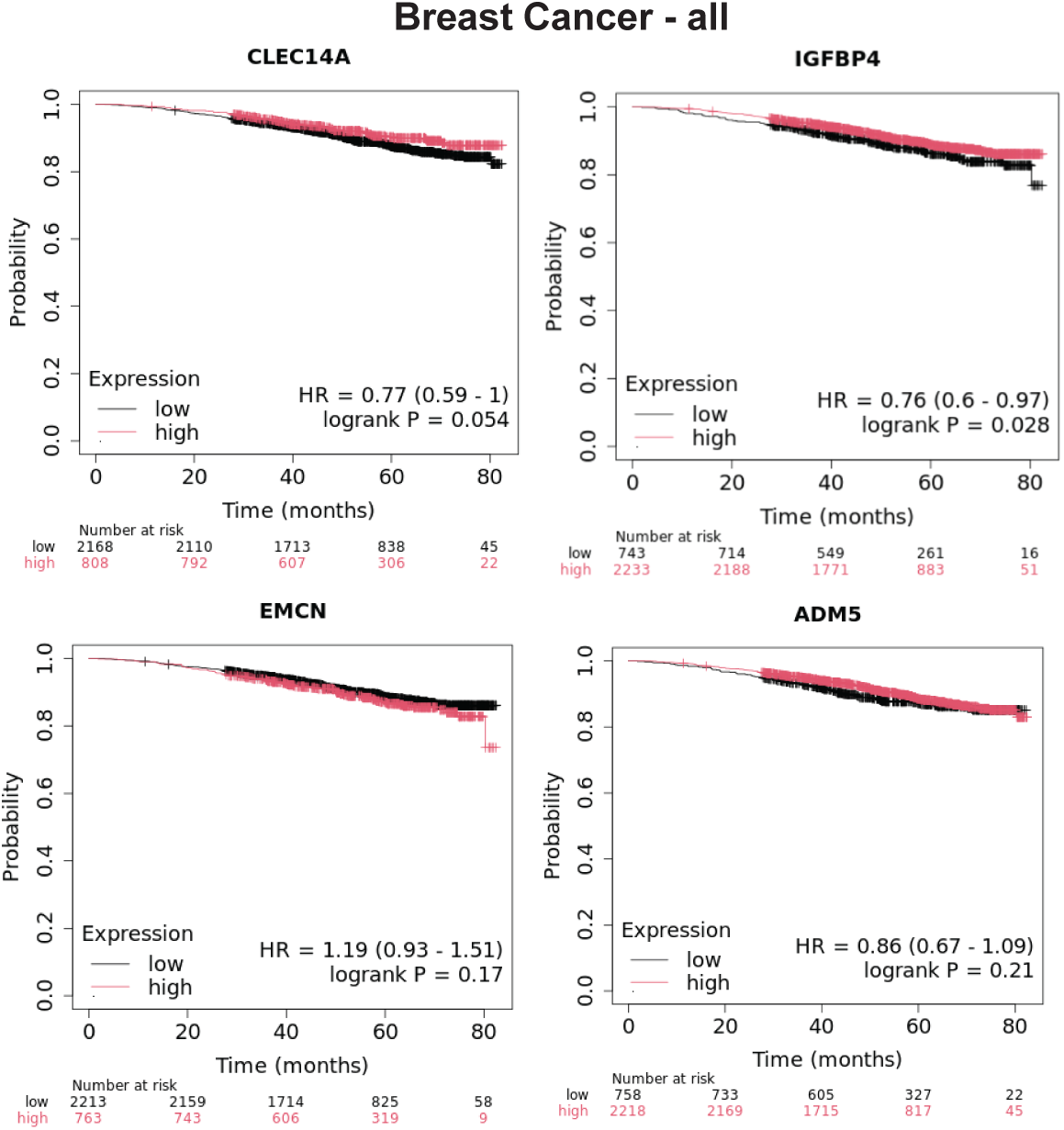
Correlation of CLEC14A, EMCN, IGFBP4, and ADM5 gene expression to overall survival in breast cancer patients (n=2976). Cox proportional hazards regression analysis was used to calculate hazard ratio (HR) h with 95% confidence intervals and log-rank p-value.

**Supplemental Figure 4.**
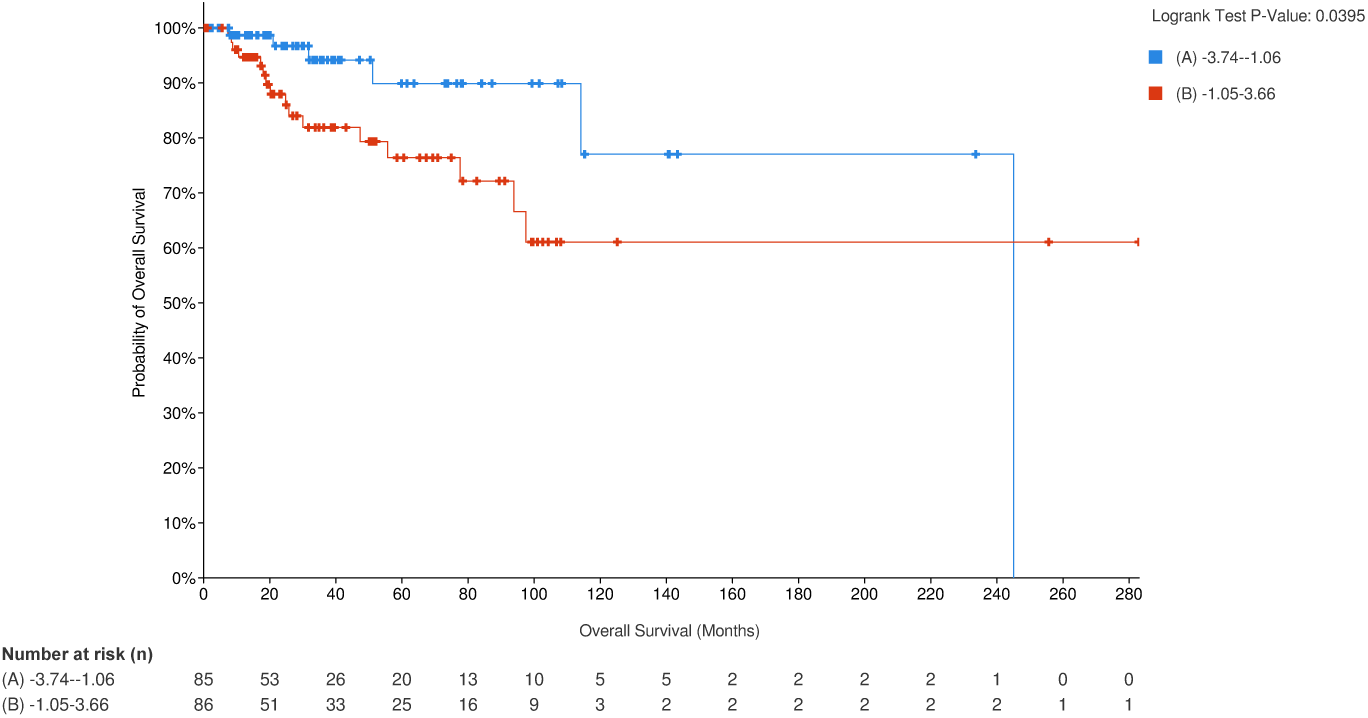
Correlation of ADM5 gene expression to overall survival in breast cancer patients with the PAM50 basal subtype (n=171) from the TCGA – BRCA dataset. Cox proportional hazards regression analysis was used to calculate hazard ratio (HR) h with 95% confidence intervals and log-rank p-value.

**Supplemental Figure 5.**
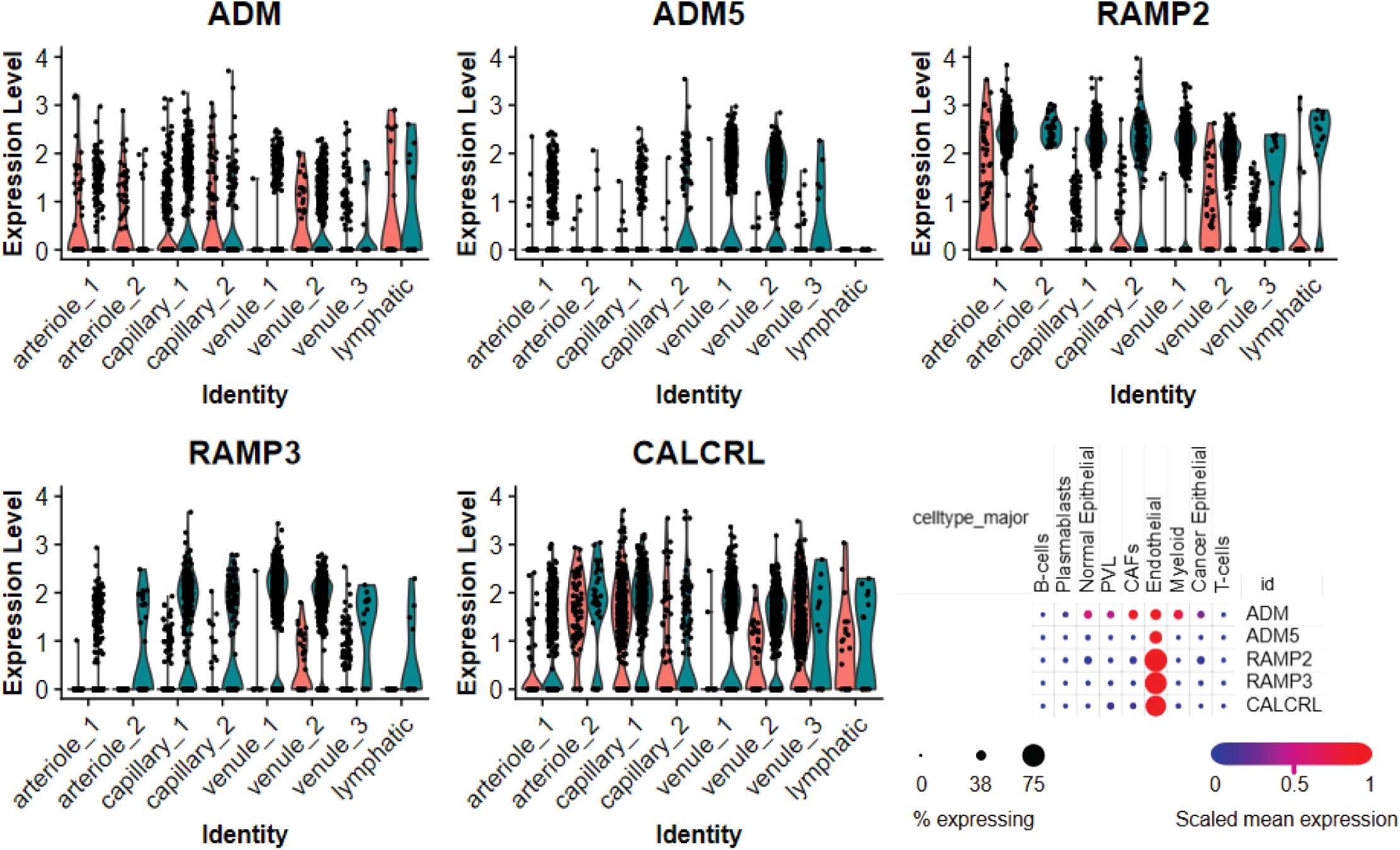
Expression of the calcitonin gene-related peptide family and related receptors. Expression of ADM, ADM5, RAMP2, RAMP3, and CALCRL. EC cluster expression split by tissue source (NEC and TEC). Cell type-specific expression of ADM, CARCRL, RAMP2, and RAMP3 in the initial Wu et al major cell type clusters. Dot color reflects gene expression level and dot size represents the percentage of cells expressing genes within each cluster.

## Notes

### Competing Interest Statement

The authors have declared no competing interest.

### Summary of Updates

These updates clarify vessel identity, strengthen the characterization of ADM5 and its signaling partners, and expand the discussion of vasodilatory pathways and endothelial specificity. We also address age-related and methodological variables, and outline future directions for in situ validation and cell-cell interaction analysis.

